# HDAC activity is dispensable for repression of cell-cycle genes by DREAM and E2F:RB complexes

**DOI:** 10.1101/2023.10.28.564489

**Authors:** Alison Barrett, Manisha R. Shingare, Andreas Rechtsteiner, Tilini U. Wijeratne, Kelsie M. Rodriguez, Seth M. Rubin, Gerd A. Müller

## Abstract

Histone deacetylases (HDACs) are pivotal in transcriptional regulation, and their dysregulation has been associated with various diseases including cancer. One of the critical roles of HDAC-containing complexes is the deacetylation of histone tails, which is canonically linked to transcriptional repression. Previous research has indicated that HDACs are recruited to cell-cycle gene promoters through the RB protein or the DREAM complex via SIN3B and that HDAC activity is essential for repressing G1/S and G2/M cell-cycle genes during cell-cycle arrest and exit.

In this study, we sought to explore the interdependence of DREAM, RB, SIN3 proteins, and HDACs in the context of cell-cycle gene repression. We found that genetic knockout of SIN3B did not lead to derepression of cell-cycle genes in non-proliferating HCT116 and C2C12 cells. A combined loss of SIN3A and SIN3B resulted in a moderate upregulation in mRNA expression of several cell-cycle genes in arrested HCT116 cells, however, these effects appeared to be independent of DREAM or RB. Furthermore, HDAC inhibition did not induce a general upregulation of RB and DREAM target gene expression in arrested transformed or non-transformed cells. Our findings provide evidence that E2F:RB and DREAM complexes can repress cell-cycle genes without reliance on HDAC activity.

## Introduction

Histone deacetylases (HDACs) play a crucial role in modulating gene expression, and functional dysregulation of their activity is linked to various medical conditions like neurodegenerative disorders, pulmonary diseases, immune disorders, and cancer (Falkenberg and Johnstone, 2014). While HDACs catalyze the deacetylation of a broad range of proteins (Shvedunova and Akhtar, 2022), they are most commonly known for removing acetyl groups from the lysines of histone tails. Hyperacetylation of histones is canonically thought to be connected to transcriptional activation, while HDAC-dependent removal of acetyl groups correlates with chromatin condensation and transcriptional repression, although recent studies have challenged this model (Morgan and Shilatifard, 2020). Aberrant expression and activity of HDACs have been found in numerous tumors (West and Johnstone, 2014), and depending on the biological context, HDACs have been shown to display both oncogenic and tumor-suppressive properties (Falkenberg and Johnstone, 2014). Three members of class I histone deacetylases – HDAC1, HDAC2, and HDAC3 – are ubiquitously expressed and are incorporated into large protein complexes (e.g. Sin3, NuRD, CoREST, SMRT, N-CoR), which get recruited to chromatin by multiple transcription factors (Emmett and Lazar, 2019; Kelly and Cowley, 2013). Several HDAC class I and panHDAC small molecule inhibitors have been approved for the treatment of hematological malignancies, such as Romidepsin (HDAC class I inhibitor, cutaneous T-cell lymphoma) and Panobinostat (panHDAC inhibitor, multiple myeloma) (Bondarev et al., 2021). HDAC inhibition (HDACi) has been shown to reduce cancer-cell proliferation and apoptosis by stimulating the expression of anti-proliferative and pro-apoptotic genes (Ramaiah et al., 2021).

A critical biological context for gene repression is the downregulation of pro-proliferative cell-cycle genes, which is required for cell-cycle arrest and exit. HDACs have been connected to the repression of two sets of these genes that control the G1/S and G2/M transitions. The timely expression of G1/S and G2/M cell-cycle genes is essential for a cell to progress through S phase, mitosis, and cytokinesis, while a loss of transcriptional repression during cell-cycle arrest and exit results in uncontrolled proliferation and oncogenic transformation (Fischer et al., 2022). In G0 and early G1, G1/S genes are repressed when E2F transcription factor binding sites within the promoters are occupied either by E2F:RB complexes, which contain an activator E2F (E2F1-3a), dimerization partner DP, and the retinoblastoma protein (RB), or by the DREAM (Dimerization partner, RB-like, E2F, And MuvB) complex, which is formed by the repressor E2Fs E2F4/5, dimerization partner DP, the pocket proteins p130/p107, and the MuvB (multi-vulval class B) core (Litovchick et al., 2007; Schmit et al., 2007). These complexes cooperate in inhibiting the expression of G1/S genes (Mages et al., 2017; Müller et al., 2017; Schade et al., 2019a; Schade et al., 2019b; Uxa et al., 2019). The expression of G2/M genes is silenced by binding of the DREAM complex to CHR promoter elements through its MuvB subunit LIN54 (Marceau et al., 2016; Müller et al., 2012; Müller et al., 2017; Müller et al., 2014). The remaining MuvB core proteins are involved in intra and inter-complex scaffolding (LIN9, LIN37, and LIN52) and histone-binding (RBBP4). LIN37 plays an important role in stabilizing the MuvB complex and positioning nucleosomes (Asthana et al., 2022; Guiley et al., 2015; Koliopoulos et al., 2022; Müller et al., 2022), and it is essential for DREAM repressor function (Mages et al., 2017; Uxa et al., 2019).

While it has been relatively well-described how E2F:RB and DREAM complexes assemble, bind to their target genes, and how these binding events correlate with gene repression, much less is known about the molecular mechanisms that prevent transcription, particularly since none of either complex’s components contain enzymatic activity. Interestingly, HDAC activity has been linked to RB- and DREAM-dependent repression (Bainor et al., 2018; Brehm et al., 1998; Ferreira et al., 1998; Ferreira et al., 2001; Lai et al., 1999; Luo et al., 1998; Magnaghi-Jaulin et al., 1998; Siddiqui et al., 2003; Zhang et al., 2000). Gene repression by RB is thought to occur through two independent mechanisms. Firstly, the multi-domain interaction between RB and E2F sterically hinders E2F’s transactivation domain from promoting transcription (Helin et al., 1993; Hiebert et al., 1992). Secondarily, RB is thought to additionally aid in G1/S gene repression by recruiting chromatin modifiers such as HDAC1/2 (Frolov and Dyson, 2004); an interaction that primarily occurs through LxCxE motifs and the RB LxCxE-binding cleft (Dick, 2007). However, since inactivating the LxCxE binding site of RB often has only limited or no effects (Andrusiak et al., 2013; Bourgo et al., 2011; Chan et al., 2001; Dahiya et al., 2000; Talluri et al., 2013; Vormer et al., 2014), it remains unclear to which extent these interactions are important for RB target gene repression.

Two models have been proposed for DREAM-dependent transcriptional repression. The most recent model stems from two structural studies and connects nucleosome positioning directly to MuvB-binding. The first of these studies showed an RBBP4/LIN37-dependent binding between MuvB and nucleosomes. DREAM stabilizes the +1 nucleosome downstream of the transcriptional start site in arrested cells, which correlates with target gene repression (Asthana et al., 2022). A second study complemented these findings by suggesting that the B-MYB-MuvB complex (MMB) may restructure nucleosome architecture during gene activation (Koliopoulos et al., 2022).

A prior model demonstrated that DREAM recruits HDAC1 through the adapter-protein SIN3B (Bainor et al., 2018). Specifically, genetic loss of SIN3B resulted in derepression of DREAM target genes in serum-starved T98G cells, and both SIN3B and HDAC1 were found to co-immunoprecipitate with DREAM. Additionally, co-immunoprecipitation of Sin3b and MuvB was detected in Rb/p107/p130 triple knockout mouse NIH3T3 cells, suggesting pocket proteins are dispensable for the interaction. An earlier report additionally showed an interaction of MuvB with SIN3B in all cell-cycle phases and proposed SIN3B to be an integral component of the MuvB core complex (Pilkinton et al., 2007). In contrast, several reports failed to detect interactions between SIN3B and DREAM/MuvB components (Adams et al., 2020; Litovchick et al., 2007; Silverstein and Ekwall, 2005; van Oevelen et al., 2008).

SIN3B and its paralog SIN3A have long been implicated in the regulation of cell-cycle genes, as well as in supporting the repression and activation of a multitude of genes in other contexts (Adams et al., 2018; Adams et al., 2020; Dannenberg et al., 2005; David et al., 2008; Pilkinton et al., 2007; Rayman et al., 2002; van Oevelen et al., 2008). Having no DNA-binding domain or enzymatic activity of their own, SIN3A and SIN3B scaffold the recruitment of chromatin modifiers like HDAC1/2, ING1/2, and KDM5A/B to histones by bridging them to transcription factors such as MAD-MAX, FOXK1, NANOG, and FAM60A (Adams et al., 2018; Chaubal and Pile, 2018). With an amino acid identity of about 60% in humans, the two SIN3 family members have both overlapping and unique functions. Generally, loss of SIN3A results in more severe phenotypes, and cells not expressing SIN3A arrest in G2/M and enter apoptosis, which is partially caused by activation of p53 (Cowley et al., 2005; Dannenberg et al., 2005). In contrast, Sin3b^-/-^ MEFs proliferate normally but show a reduced potential to arrest under growth-limiting conditions (David et al., 2008). Even though distinct SIN3A- and SIN3B-specific subcomplexes exist, both proteins have been detected at cell-cycle gene promoters in chromatin-immunoprecipitations (Bainor et al., 2018; Rayman et al., 2002; van Oevelen et al., 2010; van Oevelen et al., 2008). Combined knockdown of SIN3A and SIN3B leads to a moderate derepression of several cell-cycle genes in differentiated C2C12 cells (van Oevelen et al., 2008), suggesting that both proteins play a role in cell cycle-dependent gene regulation.

Here, we asked whether SIN3B and HDAC activity are generally required for cell-cycle gene repression and whether a loss of SIN3B would phenocopy disruption of DREAM repressor function as shown in LIN37-negative cells (Mages et al., 2017; Uxa et al., 2019). We found that while the requirement of LIN37 for DREAM repression persisted throughout a panel of cell lines, SIN3B was uniquely tied to cell-cycle gene repression in the specific context of serum-starved T98G cells. Furthermore, while a combined loss of SIN3A and SIN3B led to a moderate upregulation of cell-cycle gene mRNA expression in arrested HCT116 cells, this effect was independent of DREAM and E2F:RB repression. We further investigated the broader role of HDACs in cell-cycle gene repression and found by treating arrested cells with the small molecule HDAC inhibitors Romidepsin and Panobinostat that, while HDACs modified histones at cell-cycle genes, this activity did not generally impact transcription and is ultimately not essential to cell-cycle gene repression across cell lines. We conclude that DREAM and E2F:RB can repress cell-cycle genes independently of SIN3 proteins or HDAC activity.

## Results

### SIN3B is not essential for p53-dependent cell-cycle gene repression in HCT116 cells

Although the composition of the DREAM complex has been well described, only limited data on interaction partners that contribute to gene repression are available (Müller et al., 2022). In order to identify proteins that are enriched at the promoters of G1/S and G2/M DREAM target genes in comparison to a group of non-cell-cycle genes, we performed an *in silico* association analysis using the TFEA.ChIP tool (Puente-Santamaria et al., 2019). TFEA.ChIP utilizes the ReMap2022 database, which includes over 8000 quality-controlled ChIP-Seq datasets generated with more than 1200 chromatin-associated human proteins (Hammal et al., 2022). We selected a set of 109 G1/S and 132 G2/M genes (Suppl. Tab. 3) that we previously identified as DREAM targets (Uxa et al., 2019), and we determined which chromatin-binding proteins are enriched at the promoters of these genes in comparison to a set of 4756 genes that are not DREAM targets and that are consistently expressed throughout the cell cycle.

Indicating that the analysis is robust, G1/S genes were strongly enriched for components of E2F:RB complexes (E2F1, DP1, RB), components of DREAM (E2F4, DP1, LIN9), and repressor E2Fs (E2F6, E2F7, E2F8) (Fig. 1A). G2/M genes showed an enrichment of DREAM proteins (E2F4, DP1, LIN9), but also of B-MYB and FOXM1, which are components of the activator MuvB complexes B-MYB-MuvB and FOXM1-MuvB (Fig. 1B). Furthermore, the CCAAT-box binding proteins NFYA and NFYB were enriched at G2/M gene promoters consistent with the observation that CCAAT-boxes are often located upstream of CHR sites (Müller and Engeland, 2010). Beyond these proteins, surprisingly few other chromatin-binding factors were enriched at cell-cycle gene promoters. However, several components of histone-modifying complexes like HDAC1, HDAC2, SIN3A, SIN3B, and KDM5A were significantly enriched at both G1/S and G2/M promoters in several datasets.

**Fig. 1:**
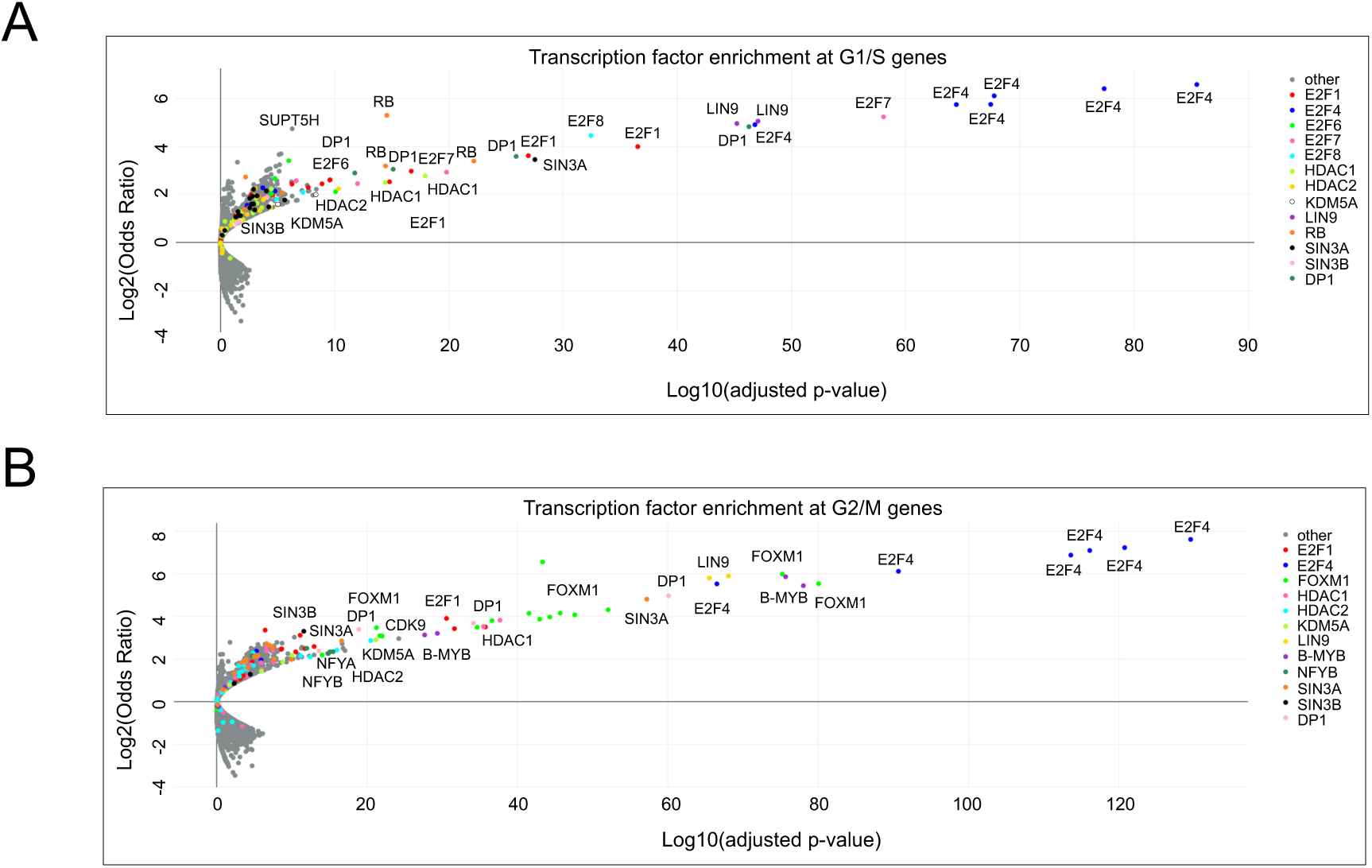
*In silico* identification of chromatin-binding proteins enriched at the promoters of G1/S and G2/M DREAM target genes. The TFEA.ChIP tool was utilized for screening the ReMap2022 database for chromatin-binding proteins enriched at the promoters of (A) G1/S (n=109) or (B) G2/M (n=132) DREAM target genes. The plots show Log2(Odds Ratio) versus Log10(adjusted p-value) for each protein (represented as a single dot) in all included ChIP-Seq experiments.

Since these proteins are all components of SIN3:HDAC complexes (Adams et al., 2018) and SIN3B has been connected to DREAM-repressor function (Bainor et al., 2018), we aimed to address whether DREAM-dependent gene repression generally relies on recruiting SIN3:HDAC, whether the loss of DREAM function upon knockout of LIN37 is phenocopied by loss of SIN3B, or whether LIN37/DREAM and SIN3B cooperate in repressing cell-cycle genes. To this end, we utilized wild-type HCT116 cells and several knockout lines (LIN37^-/-^, RB^-/-^) we had previously generated (Uxa et al., 2019) to create cells negative for SIN3B and combinations of SIN3B/LIN37 or SIN3B/RB. To minimize off-target effects, we chose a Cas9-double-nickase approach (Ran et al., 2013a) and targeted regions in exon 3 or exon 4 of the *SIN3B* gene. By probing SIN3B protein expression in clonal cell lines with two independent antibodies, we confirmed the generation of SIN3B^-/-^, SIN3B^-/-^;LIN37^-/-^, and SIN3B^-/-^;RB^-/-^ HCT116 cells (Fig. 2A).

**Fig. 2:**
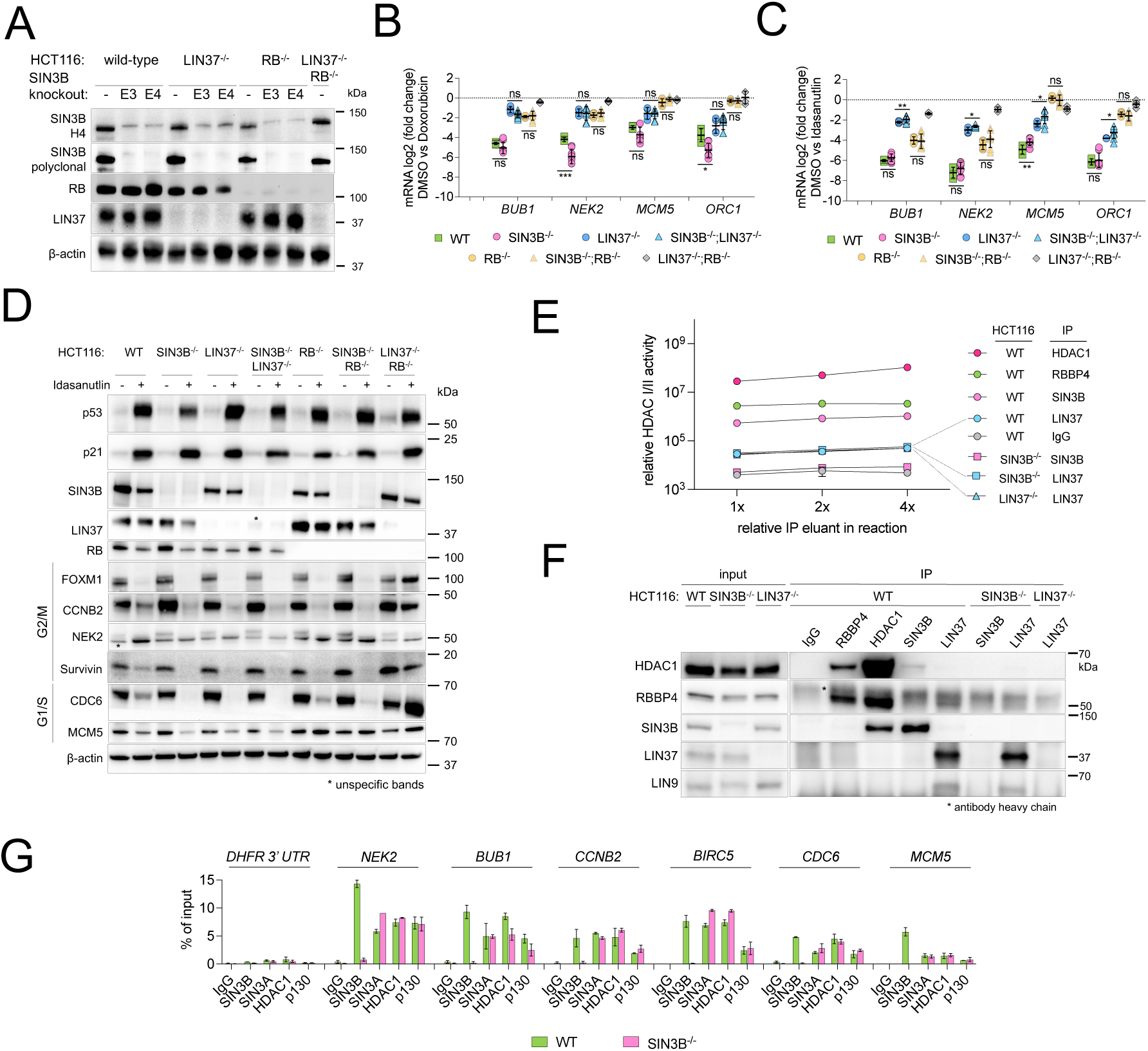
SIN3B is not essential for the repression of G1/S and G2/M cell-cycle genes as a response to DNA damage or p53 activation in HCT116 cells. (A) HCT116 cell lines negative for SIN3B were generated with a CRISPR/Cas9-nickase approach. Two pairs of guide RNAs, one targeting exon 3 and one targeting exon 4, were selected. Knockout clones were confirmed with antibodies binding epitopes within amino acids 172-228 (SIN3B-H4) or amino acids 668-758 (SIN3B polyclonal). Cells negative for SIN3B and LIN37 or RB were generated based on single knockout clones that we described earlier (Uxa et al. 2019). (B) mRNA expression of G2/M (*BUB1*, *NEK2*) and G1/S (*MCM5*, *ORC1*) cell-cycle genes was analyzed by RT-qPCR in wild-type (WT) and knockout lines after 48h treatment with 0.5 µM doxorubicin. The log2 fold change between untreated and treated cells is shown. Two independent SIN3B^-/-^, SIN3B^-/-^;LIN37^-/-^, and SIN3B^-/-^;RB^-/-^ clones were compared with wild-type cells and one LIN37^-/-^, RB^-/-^, and LIN37^-/-^;RB^-/-^ clone. The data set contains two biological replicates, and each one was measured with two technical replicates. Mean values +-SD are given. (C) Same experimental setup as in (B), but gene repression was induced by treatment with 5 µM Idasanutlin for 48 h. (D) Protein expression of HCT116 wild-type and knockout cells after treatment with DMSO or 5µM Idasanutlin for 48 h was analyzed by Western blotting. One representative experiment of three replicates is shown. (E) HDACI/II activity of samples immunoprecipitated with the indicated antibodies from HCT116 wild-type and knockout cells treated with 5 µM Idasanutlin for 48 h. Each data point contains four technical replicates of a representative experiment. Two biological replicates produced comparable results. (F) Protein expression and immunoprecipitation efficiency of the samples analyzed in (E) were evaluated by Western blotting. (G) ChIP-qPCR was performed to analyze binding of SIN3B, SIN3A, HDAC1, and the DREAM component p130 to DREAM target gene promoters in wild-type and SIN3B^-/-^cells. A non-promoter region in the 3’ untranslated region of the *DHFR* gene (*DHFR* 3’ UTR) was amplified as a negative control. Results from one out of four independent experiments are shown. Data in Figs. B, C, E, and G are presented as mean values ± SD. Significances were calculated with the two-tailed Student’s T-Test (ns – not significant, * p ≤.05, ** p ≤ .01, *** p ≤ .001). At least two biological replicates were performed for each Western Blot experiment, and results were similar.

Next, we used two different drug treatments that are known to stimulate cell-cycle arrest and cell-cycle gene repression. We induced DNA damage by treating wild-type and knockout lines with doxorubicin, or we activated the p53-p21 pathway with the MDM2 inhibitor Idasanutlin. We analyzed several representative G1/S and G2/M genes and observed a strong reduction of their expression in wild-type HCT116 cells (Fig. 2B, C). Consistent with our previously published data (Mages et al., 2017; Uxa et al., 2019), gene repression was impaired in LIN37^-/-^ and RB^-/-^ cells, and we detected a stronger loss of repression for G2/M genes in LIN37^-/-^ cells and for G1/S genes in RB^-/-^ cells. Strikingly, loss of SIN3B did not have comparable effects on cell-cycle gene repression. Upon doxorubicin treatment, repression of all analyzed cell-cycle genes did not change significantly or was even slightly, but significantly stronger in SIN3B^-/-^ than in wild-type cells. Furthermore, we did not observe any additive effects in cells negative for LIN37 or RB together with SIN3B (Fig. 2B). Cell-cycle gene expression was slightly, but mostly not significantly, elevated in SIN3B-negative cells after Idasanutlin treatment; however, loss of LIN37 or RB resulted in a much more pronounced derepression compared to SIN3B loss. Importantly, combined loss of SIN3B and RB did not reflect the almost complete loss of repression observed in LIN37^-/-^;RB^-/-^ cells (Fig. 2C).

We also analyzed several G2/M and G1/S expressed proteins in untreated and Idasanutlin-treated cells by Western blot (Fig. 2D). p53-dependent repression of G2/M expressed proteins was exclusively lost in LIN37^-/-^;RB^-/-^ cells. Deficiency for SIN3B did not result in an upregulation of G2/M protein expression, neither when it was knocked out alone nor in combination with LIN37 or RB. The G1/S proteins CDC6 and MCM5 also did not respond to loss of SIN3B; in contrast, MCM5 expression after Idasanutlin treatment was upregulated in all RB-negative cell lines (Fig. 2D).

Since it was previously reported that SIN3B serves as an adapter protein to recruit HDAC1 to the DREAM complex in T98G cells (Bainor et al., 2018), we tested whether immunoprecipitated DREAM from Idasanutlin-treated HCT116 cells contains HDAC activity. We immunoprecipitated HDAC1, SIN3B, LIN37, and RBBP4 from extracts of wild-type and knockout cells. RBBP4 is a component of MuvB as well as several chromatin-modifying complexes including the SIN3:HDAC complex (Laugesen and Helin, 2014; Müller et al., 2022). With a luciferase-based HDACI/II-activity assay, we measured robust HDAC activity in the eluates from HDAC1, SIN3B, and RBBP4 IPs (Fig. 2E). As expected, eluates immunoprecipitated with the SIN3B antibody from extracts of SIN3B^-/-^ cells showed only background activity comparable to an IgG negative control. The activity of samples immunoprecipitated with the polyclonal LIN37 antibody from wild-type extracts was slightly higher; however, HDAC activity did not change in samples precipitated from LIN37^-/-^ or SIN3B^-/-^ cells, which indicates that the antibody nonspecifically precipitates some HDAC activity independent of LIN37. These results were confirmed by Western blot analyses of the eluates (Fig. 2F). HDAC1 co-precipitated with SIN3B and RBBP4, but not with LIN37. In the LIN37 immunoprecipitations, we detected the MuvB component LIN9, but not HDAC1 or SIN3B. Thus, we did not observe endogenous DREAM and SIN3B/HDAC interactions in arrested HCT116 cells.

While several other publications also failed to show an interaction between SIN3B and MuvB complex components in immunoprecipitated samples (Adams et al., 2020; Litovchick et al., 2007; Silverstein and Ekwall, 2005; van Oevelen et al., 2008), binding of SIN3 and HDAC proteins to cell-cycle gene promoters has been shown in several cell lines by chromatin-immunoprecipitations (ChIP) (Bainor et al., 2018; Rayman et al., 2002; van Oevelen et al., 2010; van Oevelen et al., 2008) and is apparent in our meta-analysis of ChIP-seq data sets (Fig. 1). We wondered whether SIN3B, SIN3A, and HDAC1 binding to cell-cycle gene promoters in arrested HCT116 cells could be detected by ChIP. We performed ChIP-qPCR on samples from Idasanutlin-arrested wild-type and SIN3B^-/-^ cells (Fig. 2G). SIN3B was enriched at all analyzed DREAM target gene promoters in wild-type cells, and signals dropped to background level in the knockout line. Furthermore, binding of SIN3A, HDAC1, and the DREAM component p130 was detected at comparable levels in both wild-type and SIN3B^-/-^ cells.

Taken together, these data indicate that even though SIN3B binds to the promoters of DREAM target genes, it is dispensable for cell-cycle gene repression in HCT116 cells when cell-cycle arrest is induced by activation of the p53 pathway.

### Loss of SIN3B derepresses DREAM target genes in serum-starved, but not Palbociclib-treated T98G cells

Considering that we did not observe an influence of SIN3B on the repression of DREAM target genes in HCT116 cells, we next asked whether the impaired DREAM target gene repression in SIN3B^-/-^ T98G cells (Bainor et al., 2018) is phenocopied by the loss of LIN37 in the same cellular system. Our CRISPR-nickase approach for generating SIN3B and LIN37 knockouts T98G cells was less efficient than in other lines - most likely because T98G is a hyperpentaploid cell line and multiple copies of chromosome 19 that encodes for both SIN3B and LIN37 must be targeted to achieve a complete knockout. However, we were able to identify clones that did not express SIN3B, LIN37, or both proteins (Fig. 3A). Two clones of each knockout type were serum-starved, and mRNA expression of G1/S and G2/M genes was measured at several time-points after serum deprivation and compared to proliferating cells (Fig. 3B). mRNA levels of all analyzed genes were strongly reduced in starved wild-type cells. In contrast, gene repression was compromised in all knockout lines. The defect in cell-cycle gene mRNA repression in serum-starved SIN3B^-/-^ cells is consistent with previous results (Bainor et al., 2018). However, these effects were less pronounced than in LIN37^-/-^ or SIN3B^-/-^;LIN37^-/-^ cells. The derepression of cell-cycle gene expression in cells negative for SIN3B or LIN37 was also clearly detectable on the protein level (Fig. 3C).

**Fig. 3:**
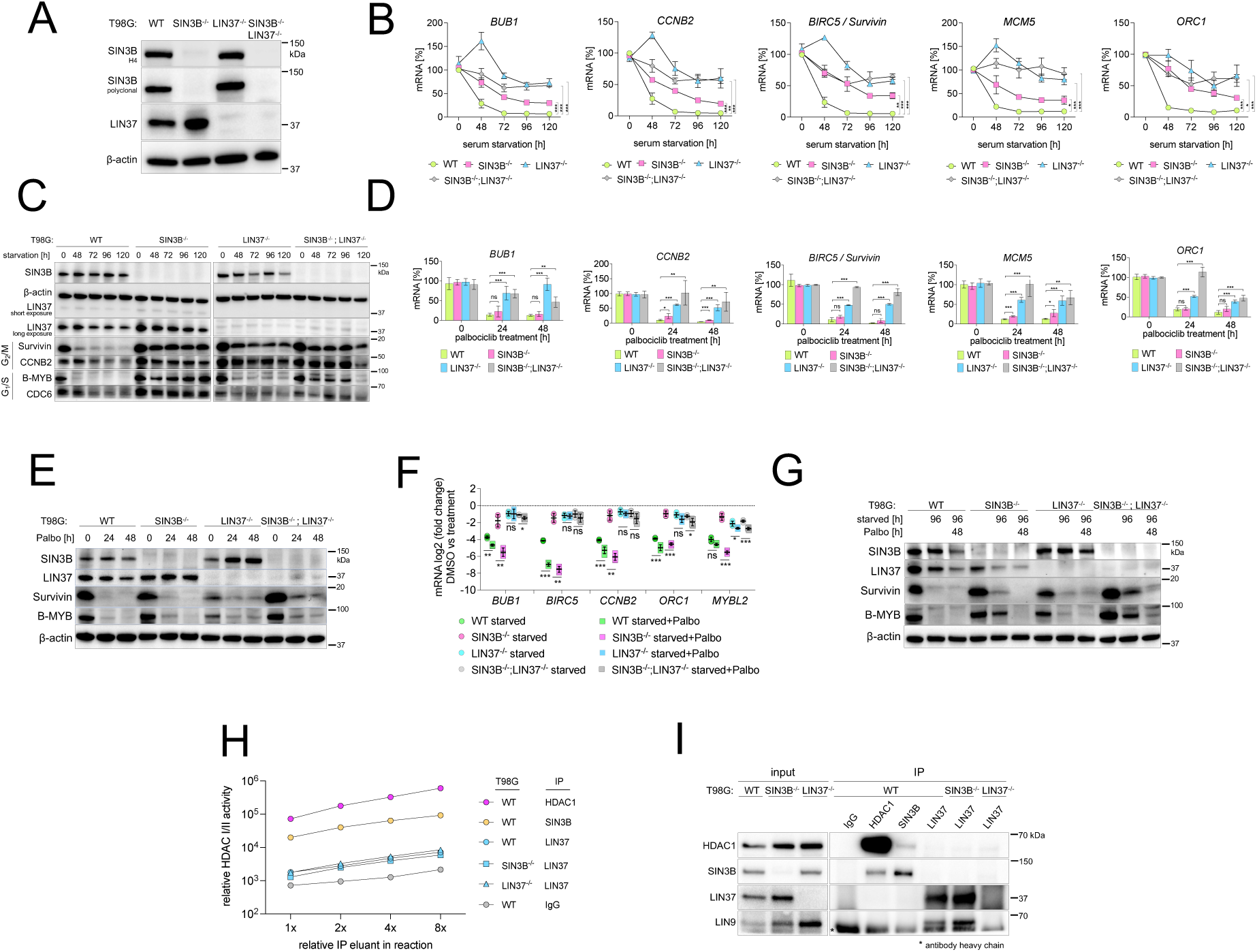
Loss of SIN3B does not phenocopy LIN37 deficiency in T98G cells. (A) A CRISPR/Cas9-nickase approach to introduce mutations in exon 4 of SIN3B and exon 6 of LIN37 was applied to generate cell lines negative for SIN3B, LIN37, or both proteins. SIN3B knockout clones were confirmed with antibodies targeting amino acids 172-228 (SIN3B-H4) or amino acids 668-758 (SIN3B polyclonal). LIN37 knockout was confirmed with a polyclonal antibody raised against full-length LIN37. (B) mRNA expression of G2/M (*BUB1*, *CCNB2*, *BIRC5*) and G1/S (*MCM5, ORC1*) cell-cycle genes was analyzed by RT-qPCR in wild-type and knockout lines arrested by serum-starvation. Two independent SIN3B^-/-^, LIN37^-/-^, and SIN3B^-/-^;LIN37^-/-^ clones were compared with two wild-type (WT) clones measured with two technical replicates each. Significances of differences in gene expression after 120 h of serum deprivation were analyzed with the two-tailed Student’s T-Test. (C) Protein expression of one of the wild-type and knockout clones measured in (C) was analyzed by Western blotting. Similarly, (D) mRNA expression and (E) protein levels of cell-cycle genes were analyzed in two wild-type or knockout lines treated with 10 µM Palbociclib for 24 or 48 hours. (F) Indicated wild-type and knockout lines (two clones each) were serum-starved for 96hrs with or without 10 µM Palbociclib for the final 48 h. mRNA was measured (two technical replicates each) and compared with untreated wild-type mRNA levels. (G) Samples shown in (F) were additionally analyzed for protein expression by Western blotting. (H) HDACI/II activity of samples immunoprecipitated from T98G wild-type and knockout cells serum-starved for 96 h with the indicated antibodies. Each data point contains four technical replicates of a representative experiment. Two biological replicates produced similar results. (I) Protein expression and immunoprecipitation efficiency of the samples analyzed in (H) were evaluated by Western blotting. Data in Figs. B, D, F, and H are given as mean values ± SD, and significances were calculated with the two-tailed Student’s T-Test (ns – not significant, * p ≤.05, ** p ≤ .01, *** p ≤ .001). At least two biological replicates were performed for each Western Blot experiment, and results were similar.

To analyze cell-cycle gene repression in a setting other than serum deprivation, we treated the T98G lines with Palbociclib. We chose Palbociclib (as opposed to Idasanutlin) to directly inhibit CDK4/6 since T98G cells do not express wild-type p53. Surprisingly, while the observed highly significant increase in cell-cycle gene expression detected in starved LIN37^-/-^ or SIN3B^-/-^;LIN37^-/-^ persisted in Palbociclib-treated cells, we did not observe a robust loss of mRNA repression in SIN3B^-/-^ cells after 24 or 48 hours of treatment (Fig. 3D). Additionally, Palbociclib treatment led to comparable repression of DREAM targets in wild-type and SIN3B^-/-^ cells on the protein level, while increased expression was detected in LIN37^-/-^ and SIN3B^-/-^;LIN37^-/-^ cells (Fig. 3E). We next asked whether addition of Palbociclib reinforces cell-cycle gene repression in T98G knockout lines that are serum-starved. To this end, we compared cell-cycle gene expression on mRNA and protein levels in cells that were either serum-starved for 96h or starved for the same period but with the addition of Palbociclib for the final 48 hours. As observed before (Fig. 3B, C), loss of SIN3B or LIN37 resulted in an elevated expression of cell-cycle genes in starved cells compared to the parental line (Fig. 3F, G). The addition of Palbociclib increased the repression in the wild-type cells, and the measured genes were repressed to the same extent or even stronger in the SIN3B knockouts. In contrast, addition of Palbociclib to LIN37-negative serum-starved cells led only to minimal changes in cell-cycle gene expression. Comparable effects were also observed on the protein level (Fig. 3F, G).

Taken together, our data demonstrate that the reduction in cell-cycle gene repression in serum-starved SIN3B^-/-^ T98G cells can be bypassed by directly inhibiting CDKs, but only in LIN37-positive cells that can assemble a functional DREAM complex. We propose that the observed defect in serum-starved SIN3B^-/-^ T98G cells is not caused by a loss of DREAM repressor function, but by upstream mechanisms that result in an impaired CDK inhibition, which prevents the formation of DREAM and RB/E2F complexes.

To analyze whether endogenous DREAM contains HDAC activity in T98G cells, we immunoprecipitated HDAC1, SIN3B, and LIN37 from serum-starved T98G cells and performed HDAC activity assays with the eluates. As expected, we detected strong HDAC activity in the samples containing purified HDAC1 and SIN3B (Fig. 3H). The HDAC activities in eluates precipitated with the LIN37 antibody were comparable between samples obtained from LIN37 positive and negative cell lines suggesting that the signals are nonspecific. The data obtained from HDAC assays are in line with Western blot results that show a coprecipitation of HDAC1-SIN3B and LIN37-LIN9, but no interaction of MuvB components with SIN3B or HDAC1 (Fig. 3I). As we observed in HCT116 cells (Fig. 2E, F), we did not find evidence of proteins containing HDAC activity interacting with DREAM in starved T98G cells.

### Sin3B is not essential for cell-cycle gene repression in arrested C2C12 cells

Given that we and others (Bainor et al., 2018) observed defects in cell-cycle gene repression in serum-deprived SIN3B^-/-^ T98G cells, we wondered whether this effect generally occurs during serum-starvation. Since HCT116 cells cannot efficiently be arrested by serum deprivation and rapidly induce apoptosis following serum withdrawal (Blagosklonny et al., 1997), we created Sin3b-negative mouse C2C12 cells (Fig. 4A) and compared cell-cycle gene repression after serum-starvation with wild-type and Lin37^-/-^ cells (Fig. 4B) (Mages et al., 2017; Uxa et al., 2019).

**Fig. 4:**
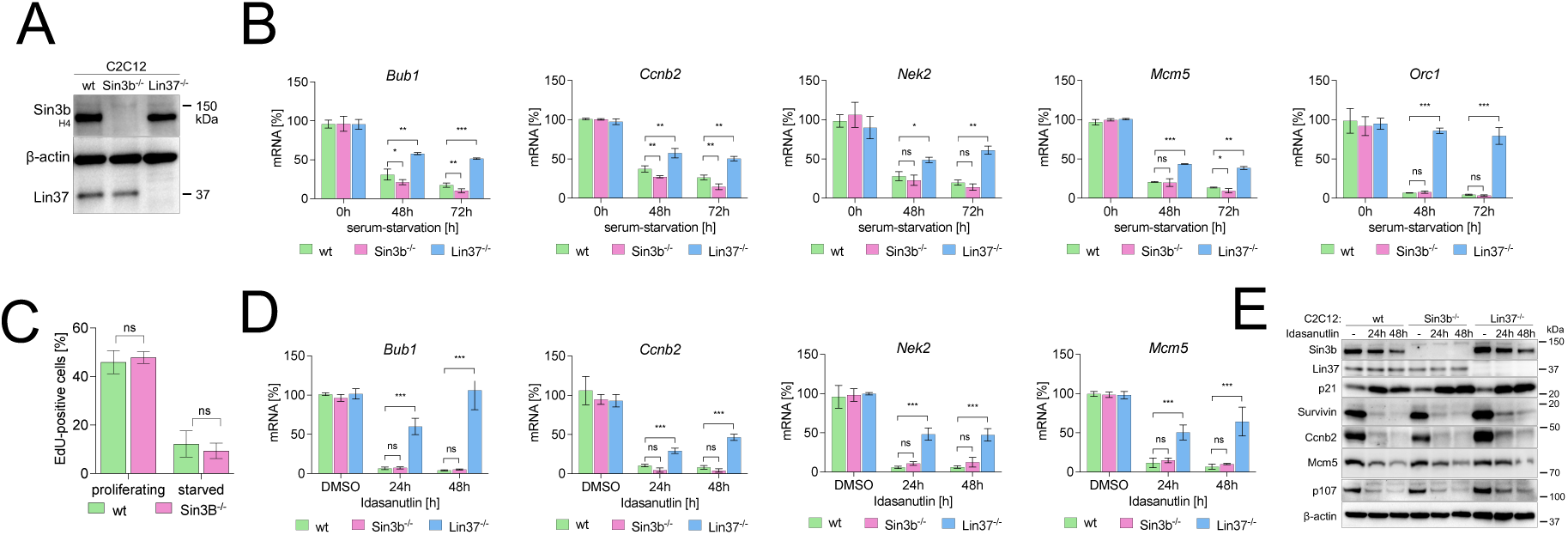
Sin3b knockout does not phenocopy loss of Lin37 in mouse C2C12 cells. (A) A CRISPR/Cas9-nickase approach was applied to generate cell lines negative for Sin3b. Sin3b knockout clones were confirmed with antibodies targeting an epitope within amino acids 172-228 (SIN3B-H4). Lin37^-/-^ C2C12 cells were described before (Mages et al. 2017). (B) mRNA expression of cell-cycle genes was analyzed by RT-qPCR in wild-type and knockout lines arrested by serum starvation over 48 and 72h. Two wild-type, one Lin37^-/-^, and three SIN3B^-/-^ clones were measured, with two technical replicates each. (C) EdU incorporation was analyzed in wild-type and Sin3B^-/-^ lines (3 clones each) after 48 h of serum deprivation. (D) mRNA expression of cell-cycle genes was analyzed by RT-qPCR in the same wild-type and knockout lines shown in (B) treated with 5 µM Idasanutlin for 24 and 48 h. (E) Protein expression of one clone analyzed in (D) was studied by Western blot. Data in (B), (C), and (D) Data are presented as mean values ± SD. Significances were calculated with the two-tailed. Student’s T-Test (ns – not significant, * p ≤.05, ** p ≤ .01, *** p ≤ .001). Western Blot experiment were performed with at least two biological replicates with similar results.

Starvation for 48 or 72 hours led to repression of G1/S and G2/M genes in the wild-type cells. Loss of Sin3b did not result in defective repression of these genes, but rather, the Sin3b^-/-^ cells appeared to have slightly stronger repressive trends than wild-type cells. In contrast, all measured genes were significantly derepressed in serum-starved Lin37^-/-^ C2C12 cells (Fig. 4B). Since it had been previously shown that Sin3b^-/-^ MEFs exit the cell cycle less efficiently than wild-type cells when serum-starved (David et al., 2008), we tested EdU incorporation in proliferating and starved wild-type and Sin3B^-/-^ cells, but we did not find significant differences (Fig. 4C). Idasanutlin-treatment also reduced cell-cycle gene mRNA (Fig. 4D) and protein (Fig. 4E) expression similarly in wild-type and Sin3b^-/-^ cells and to a lesser extent in Lin37^-/-^. Based on these findings, we conclude that loss of SIN3B does not generally influence the response of cells to the withdrawal of mitogenic stimuli.

### Combined loss of SIN3A and SIN3B increases the mRNA expression of multiple cell-cycle genes in arrested cells independently of DREAM or RB

Since we did not find deregulation of DREAM targets in arrested SIN3B^-/-^ HCT116 cells, we asked whether SIN3A can compensate for loss of SIN3B in these cells. It has been demonstrated that depletion of SIN3A results in cell-cycle arrest and apoptosis through activation of CDKN1A/p21 in a p53-dependent and -independent manner (Dannenberg et al., 2005). Based on these data, we refrained from knocking out SIN3A and instead chose a siRNA-based approach to reduce SIN3A expression. First, we tested the knockdown efficiency of four independent SIN3A siRNAs in proliferating HCT116 cells. All four siRNAs drastically reduced the protein expression of SIN3A, while SIN3B levels were increased (Suppl. Fig. 1A). Even though SIN3A was reduced below the detection level with all four siRNAs, accumulation of p53 and p21 only occurred with siRNAs 1, 2, and 4, suggesting that sufficient SIN3A remained in cells treated with siRNA 3. Cell-cycle protein expression behaved inversely to p53 and p21 levels: mitotic and S phase regulators were repressed upon transfection of SIN3A siRNAs 1, 2, and 4 (Suppl. Fig. 1A). The observed repression of cell-cycle genes was be recapitulated on the mRNA level (Suppl. Fig. 1B).

Next, we analyzed whether SIN3A knockdown in Idasanutlin-treated wild-type and SIN3B^-/-^ cells influenced repression of cell-cycle genes. SIN3A protein expression was already reduced in arrested HCT116 cells without siRNA treatment, and RNA interference decreased SIN3A levels still further. Protein expression of G2/M and G1/S cell-cycle regulators was strongly repressed upon Idasanutlin treatment in both wild-type and SIN3B^-/-^ cells, and knockdown of SIN3A did not result in a detectable upregulation (Suppl. Fig. 1C). Knockdown of SIN3A in wild-type HCT116 cells led to minor effects regarding the mRNA expression of the analyzed cell-cycle genes, while a combined loss of SIN3B and SIN3A resulted in an upregulation throughout all analyzed genes (Suppl. Fig. 1D). Next, we performed transcriptome analyses to identify genes deregulated in Idasanutlin-arrested WT, SIN3B^-/-^, SIN3A knockdown, and SIN3B^-/-^ + SIN3A knockdown HCT116 cells. We found comparable numbers of genes significantly (p<0.05) up- and downregulated (≥1.5fold) within all three conditions when compared to wild-type cells (Fig. 5A, Suppl. Tab. S4). Out of 268 LIN37/DREAM target genes we had identified earlier (Uxa et al., 2019), only 3 were upregulated in non-proliferating SIN3B^-/-^ cells, and 7 in non-proliferating SIN3A knockdown cells. However, this number increased to 57 (21%) in cells depleted of SIN3A and SIN3B (Fig. 5A, B, Suppl. Tab. S4). GO analyses confirmed that upregulated genes connected to cell-cycle relevant processes were only enriched in SIN3A/B-depleted cells (Fig. 5C). Interestingly, knockdown of SIN3A resulted in upregulation of genes sets connected to cilium organization and assembly, while genes upregulated in SIN3B^-/-^ cells only produced 7 predominantly broad terms associated with high False Discovery Rate (FDR) values.

**Fig. 5:**
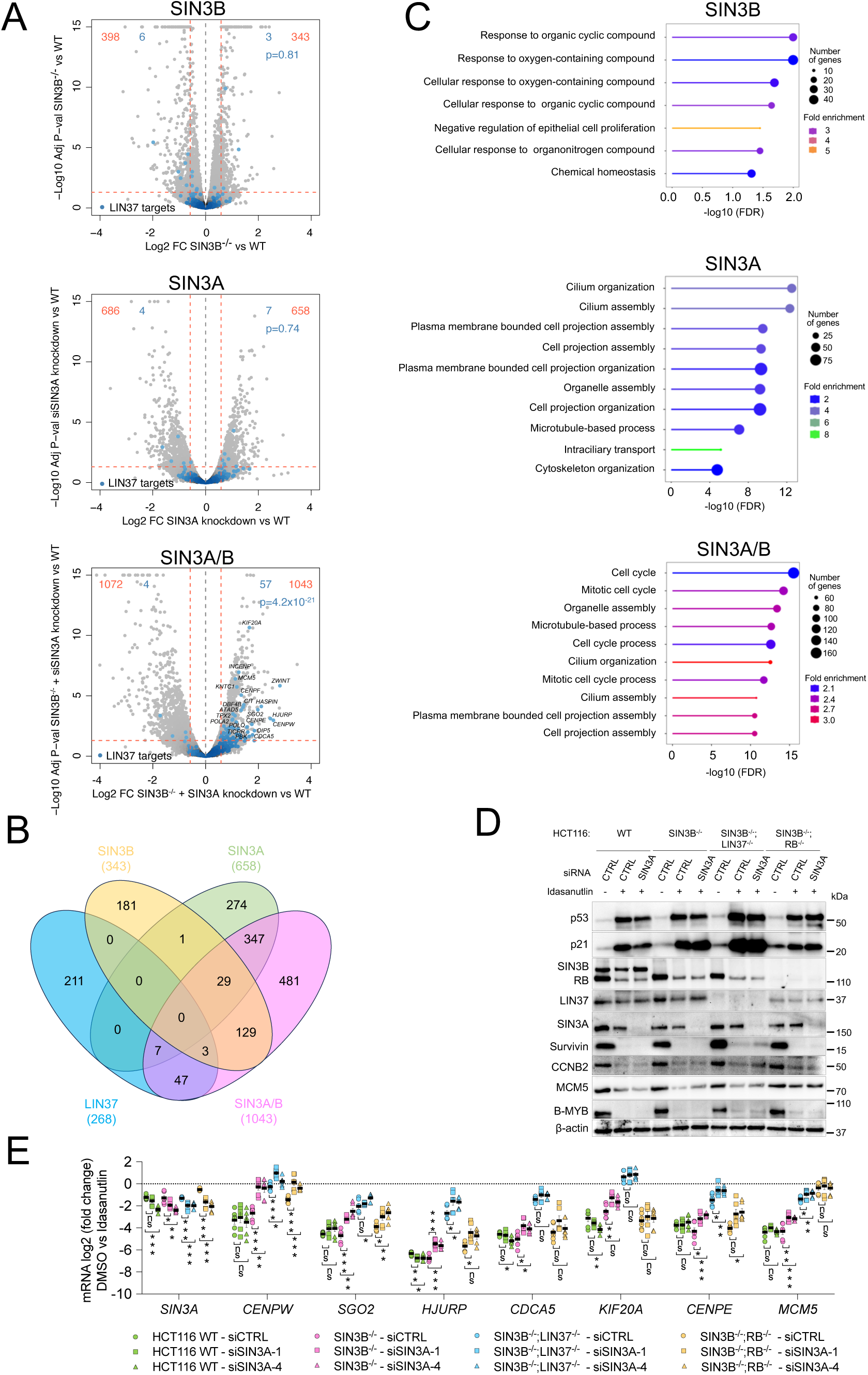
Combined depletion of SIN3A and SIN3B derepresses a subset of cell-cycle genes independently of DREAM or RB. Transcriptome analyses were performed with HCT116 wild-type (WT) and SIN3B^-/-^ HCT116 cells transfected either with a non-targeting siRNA or with SIN3A siRNAs for 48 h and treated with Idasanutlin for the final 24 h. (A) Volcano plots showing up and downregulated genes in comparison to wild-type cells. Numbers of significantly regulated genes (p<0.05) with an FC ≥1.5 are shown in red. Genes identified as LIN37/DREAM targets before (Uxa et al., 2019) are highlighted in blue. The p-values indicate the probability that the respective overlap between LIN37 and SIN3-regulated genes could be observed by chance (hypergeometric test). (B) The number and overlap of LIN37 target genes identified in Uxa et al. and genes upregulated (p<0.05; FC ≥1.5) in Idasanutlin-treated HCT116 cells depleted of SIN3A, SIN3B, or both proteins. (C) GO analyses (biological processes) of significantly upregulated (p<0.05; FC ≥1.5) genes. The top ten hits based on their false discovery rate (FDR) are shown. (D) Wild-type, SIN3B^-/-^, SIN3B^-/-^;LIN37^-/-^, and SIN3B^-/-^;RB^-/-^ knockout lines were transfected with non-targeting or SIN3A siRNAs for 48 h, and Idasanutlin was applied for the final 24 h. Protein levels were analyzed by Western blotting. A representative blot of cells transfected with either a non-silencing control RNA (CTRL) or SIN3A siRNA 1 is shown. A biological replicate with SIN3A siRNA 4 produced similar results. (E) mRNA levels of genes identified as significantly upregulated in SIN3A/B-depleted cells in the transcriptome analysis were evaluated by RT-qPCR. Cells were treated as described in (D), but additionally transfected with SIN3A siRNA 4. Averages (mean values ± SEM) of three biological and two technical replicates are given. Significances were calculated with the two-tailed Student’s T-Test (ns – not significant, * p ≤.05, ** p ≤ .01, *** p ≤ .001).

We then aimed to analyze whether the observed cell-cycle gene upregulation depended on DREAM or RB. We transfected wild-type, SIN3B^-/-^, SIN3B^-/-^;LIN37^-/-^, and SIN3B^-/-^;RB^-/-^ lines with non-targeting or SIN3A siRNAs to analyze the expression of several cell-cycle genes that were identified as upregulated after SIN3A/B depletion in the RNA-Seq experiment by RT-qPCR. Knockouts and p53 induction were confirmed by Western blotting (Fig. 5D). While derepression of the *CENPW* and *SGO2* genes in Idasanutlin-treated SIN3A/B-depleted cells was comparable to the effects observed in LIN37-negative cells, repression of the other analyzed genes relied more strongly on LIN37/DREAM or RB than on SIN3A/B (Fig. 5E). Furthermore, we observed that the derepression caused by loss of LIN37 or RB was generally further increased after depletion of SIN3A/B. These additive effects suggest that SIN3A/B repress cell-cycle genes independent of DREAM and RB. Comparable trends could also be observed by analyzing the expression of *BUB1*, *NEK2*, and *ORC1* (Suppl. Fig. 1D, E), which are cell-cycle genes that were found to be significantly derepressed after SIN3A/B depletion in our RNA-seq experiment. Thus, it is likely that SIN3A/B contribute to the repression of more cell-cycle genes than we identified in the RNA-seq screen in a DREAM and RB-independent manner. The moderate upregulation of mRNA expression observed for several cell-cycle genes after loss of SIN3A/B did not translate to detectable changes on protein level (Fig. 5D).

Taken together, loss of SIN3B or SIN3A alone did not lead to an upregulation of cell-cycle gene expression in arrested HCT116 cells. Combined depletion moderately increased cell-cycle gene mRNA expression, but this effect appeared to be independent of DREAM and RB. Moreover, effects from combined SIN3B and SIN3A depletion were minor compared to the derepression observed after loss of LIN37 or RB.

### Inhibition of HDAC activity does not broadly upregulate cell-cycle genes in arrested cells

Since we detected binding of HDAC1 to cell-cycle gene promoters in HCT116 cells independent of SIN3B (Fig. 2G), we asked whether histone tail acetylation at cell-cycle gene promoters changes during cell-cycle arrest and HDACi, and whether the repression of G2/M and G1/S genes in arrested cell lines is reduced when HDAC activity is inhibited. Since HDAC1/2 inhibition itself results in the upregulation of cell-cycle inhibitors like p21 and induces cell-cycle arrest (Vinodhkumar et al., 2008; Yamaguchi et al., 2010; Zupkovitz et al., 2010), we arrested HCT116 cells with Idasanutlin first and then added the HDAC1/2 inhibitor Romidepsin (Nakajima et al., 1998; Ueda et al., 1994). ChIP-qPCR analyses showed that H3K27 acetylation at the promoters of several MuvB target genes was reduced upon Idasanutlin treatment, and addition of Romidepsin reversed this effect. In contrast, H3K27 trimethylation was strongly reduced in Romidepsin-treated cells (Fig. 6A). Next, we measured the expression of 19 representative G2/M and 13 G1/S genes and compared their repression in cells treated exclusively with Idasanutlin or with Idasanutlin and Romidepsin (Fig. 6B). For both groups of genes, we did not observe a significant loss of the average repression upon HDAC1/2 inhibition, although several genes like SGO2, *NEK2*, *E2F8*, *RBL1*, and *ORC1* were slightly, but significantly upregulated. Furthermore, except for *SGO2*, expression of genes that were upregulated in SIN3A/B-depleted cells did not increase upon HDACi. In contrast, a set of genes that had been previously reported to be upregulated in proliferating HCT116 cells after HDACi (LaBonte et al., 2009) showed a highly significant average increase in expression (Fig. 6B). Further demonstrating the efficacy of the drug treatments, Western blot analysis confirmed upregulation of p53 and p21 in response to Idasanutlin treatment and showed a strong increase of acetylated histone H3 upon Romidepsin treatment (Fig. 6C). Expression of G2/M and G1/S proteins was strongly repressed in Idasanutlin-treated cells, and addition of Romidepsin did not increase protein levels (Fig. 6C). While the observed reduction in H3K27ac levels correlates with Idasanutlin-induced gene repression, the increase that follows HDACi but not cell-cycle arrest (Fig. 6A) does not result in an upregulation of gene expression. These data indicate that cell-cycle gene repression can be maintained even when the chromatin at the promoters shows hallmarks of actively expressed genes.

**Fig. 6:**
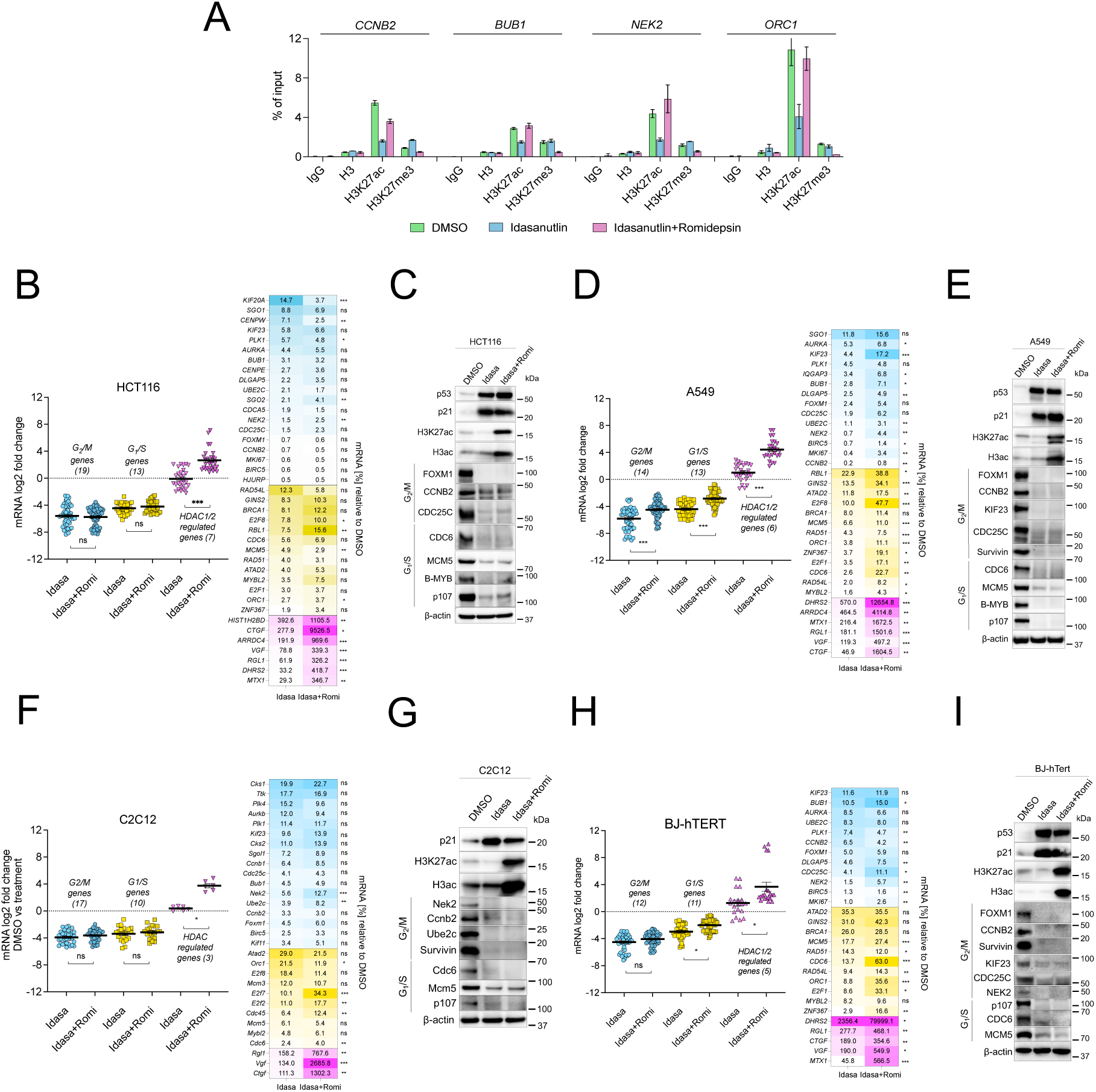
HDAC activity is not generally required for cell-cycle gene repression in arrested cells. In all the following experiments, cells were treated with 5 µM Idasanutlin for 48 h and 4 nM Romidepsin for the final 24 h. (A) Histone modifications at cell-cycle gene promoters in HCT116 cells were analyzed by ChIP-qPCR. mRNA expression was evaluated by RT-qPCR and compared to DMSO-treated cells for each respective line. One representative experiment with two technical replicates (mean values ± SD) is shown. A biological replicate produced similar results. Data are presented as gene-set clusters (left) and individual genes (right) in: (B) HCT116, (D) A549, (F) C2C12, and (H) BJ-hTERT cells. The datasets contain two biological replicates with two technical replicates each. Mean values ± SEM are shown, and significances were calculated with the two-tailed Students T-Test (ns – not significant, * p ≤.05, ** p ≤ .01, *** p ≤ .001). Protein expression and histone acetylation were evaluated via Western blotting with the indicated antibodies for: (C) HCT116, (E) A549, (G) C2C12, and (I) BJ-hTERT cells. A biological replicate for each Western blot experiment produced similar results.

To analyze whether additional HDACs that are not inhibited by Romidepsin influence the repression of MuvB target genes, we repeated the experiment with the pan-HDAC inhibitor Panobinostat (George et al., 2005) and obtained comparable results (Suppl. Fig 2A, B). To test the effect of HDACi in additional cell lines, we treated A549 lung carcinoma cells with Idasanutlin and Romidepsin. In these cells, multiple G2/M and G1/S genes were significantly upregulated in Idasanutlin-treated cells after Romidepsin treatment, even though the average increase of expression was lower than the set of HDAC-dependent control genes (Fig. 6D). However, this upregulation of mRNA level did not lead to a detectable increase in protein expression (Fig. 6E). Treatment of Idasanutlin-arrested A549 cells with the pan-HDAC inhibitor Panobinostat led to some minor but predominantly non-significant changes in mRNA expression of G2/M and G1/S genes (Suppl. Fig. 2C), and Panobinostat-treatment also did not translate to detectable changes in protein expression (Suppl. Fig. 2D). We next analyzed effects of HDAC1/2 inhibition in arrested non-transformed mouse C2C12 cells. Comparable to HCT116 cells, mRNA expression of HDAC-dependent genes was strongly increased, while levels of G1/S and G2/M gene mRNAs were on average not significantly upregulated (Fig. 6F). Even though several genes were significantly upregulated (Fig. 6F), these changes did not translate to the protein level (Fig. 6G). To analyze the effects of HDACi in a human, non-transformed cell line, we repeated the experiment BJ-hTert cells. Addition of Romidepsin to Idasanutlin-treated cells slightly increased the expression of several G2/M genes (*CDC25C*, *NEK2*, *MKI67*), while the expression of others did not change significantly (*AURKA*, *UBE2C*, *FOXM1*) or was further reduced (*PLK1*, *CCNB2*). The set of analyzed G1/S genes generally appeared to be more strongly influenced by Romidepsin treatment, particularly *CDC6*, *ORC1*, *E2F1*, and *ZNF367*, which were significantly upregulated. However, several of the other analyzed G1/S genes did not respond to HDACi (*ATAD2*, *BRCA2*, *RAD51*, *MYBL2*), while all control genes were significantly upregulated (Fig. 6H). Again, no change in repression of cell-cycle gene protein expression could be detected by Western blotting (Fig. 6H).

Finally, we asked whether HDACi in Palbociclib-treated T98G wild-type and knockout cells results in defects in cell-cycle gene repression since DREAM target genes can be repressed in SIN3B^-/-^ cells in this context (Fig. 3). While the mRNA expression of the HDAC-dependent control gene *CTGF* was significantly upregulated in all tested clones, HDACi by Romidepsin or Panobinostat did not reduce Palbociclib-dependent repression of *BUB1* and *ORC1*. In contrast, repression of these genes was enforced in wild-type and SIN3B^-/-^ cells, which most likely occurred through upregulation of p21, CDK1/2 inhibition, and an increased formation of DREAM since this effect was not observed in LIN37^-/-^ cells (Suppl. Fig. 2E).

Taken together, our results provide evidence that HDACi in arrested transformed and non-transformed cells does not generally derepress DREAM and E2F:RB target genes. Interestingly, we see significant derepression of some cell-cycle genes, however the affected genes are variable between cell lines and do not affect protein levels. We conclude that HDAC involvement in cell-cycle gene regulation is not a general mechanism of repression by DREAM and E2F:RB complexes.

## Discussion

Cell-cycle dependent gene regulation has been studied extensively for decades, but the mechanisms by which MuvB and E2F:RB complexes mediate the precisely timed repression or activation of several hundred genes remain poorly understood. In this study, we aimed to analyze to what extent HDAC activity contributes to the repression of cell-cycle genes by DREAM and E2F:RB complexes. The involvement of HDAC-containing complexes in the regulation of cell-cycle genes has been controversially discussed. The earliest reports proposing the involvement of HDACs in the repression of E2F target genes showed a direct LxCxE-dependent binding of HDAC1 to RB and suggested a dependence of RB transcriptional repressor function on HDAC1 based on promoter reporter assays (Brehm et al., 1998; Ferreira et al., 1998; Magnaghi-Jaulin et al., 1998) or RT-PCR (Luo et al., 1998). Interestingly, one study observed that RB-dependent repression of some of the analyzed G1/S genes relied on HDAC activity, while others were resistant (Luo et al., 1998). In addition to HDAC1, more than 100 proteins have been reported to interact with RB (Morris and Dyson, 2001), and a subset of them has been validated to use a LxCxE motif for binding (Dick, 2007). Given that HDAC1 has a low affinity to RB in comparison to other partners that potentially compete for LxCxE-dependent RB binding (Putta et al., 2022), it remains unclear to what extent HDAC contributes to RB repressor function in particular biological settings. Furthermore, several studies provided evidence that, depending on the context, mutation of the RB LxCxE binding cleft that disrupts the multitude of potential binding interactions has only limited and in some cases no effects on overall cell-cycle gene repression, cell-cycle arrest or exit, and induction of carcinogenesis (Andrusiak et al., 2013; Bourgo et al., 2011; Dahiya et al., 2000; Talluri et al., 2013; Vormer et al., 2014). The further complexity of this system was established when the p130/p107-containing mammalian DREAM complex was discovered (Litovchick et al., 2007; Schmit et al., 2007). DREAM is the main transcriptional repressor of G2/M genes at CHR promoter elements, and it represses G1/S genes together with E2F:RB complexes at E2F promoter sites (Fischer and Müller, 2017). HDAC1 had been shown to bind p130/p107 and RB equally well and in a LxCxE-dependent manner both *in vitro* and when overexpressed in cells (Ferreira et al., 1998). However, we later demonstrated that p130/p107 are incorporated into the DREAM repressor complex via a LxCxE-dependent interaction with LIN52 (Guiley et al., 2015), which makes recruitment of additional proteins through the LxCxE binding cleft unlikely. This example highlights that even though an interaction can be detected in a specific experimental setup, it may not be relevant in a particular physiological context.

Given that none of the DREAM components contain any enzymatic activity and that p130/p107 cannot bind chromatin modifiers via its LxCxE binding cleft when present in DREAM, the question arises if other DREAM components can recruit such proteins. The obvious candidate for such a mechanism is the MuvB core component RBBP4, which is also found in several chromatin-modifying complexes such as PRC2, NuRD, CoRest, and SIN3 (Laugesen and Helin, 2014). However, evidence regarding binding of histone modifiers to DREAM components is limited. While IP/MudPIT analysis after immunoprecipitating tagged p130, LIN9, LIN37, or LIN52 from T98G cells identified all DREAM components, no other interactors were detected in all samples (Litovchick et al., 2007). Particularly, no SIN3B or HDAC1/2 were detected in any of the samples, even though peptides of HDAC3 were identified in LIN54 IPs and peptides of SIN3A in LIN37 IPs (Litovchick et al., 2007). In contrast, tandem mass-spectrometry of samples gained by immunoprecipitating overexpressed SIN3B from immortalized MEFs identified all DREAM and MMB components except LIN52 as binding partners (Bainor et al., 2018). The strong discrepancies in the results of both studies could be caused by the expression and/or precipitation of the different proteins. However, in a recent study that extensively analyzed the interaction network of SIN3A and SIN3B, no proteins specific for MuvB complexes were found in samples precipitated with tagged SIN3A/B from HEK extracts except RBBP4 (Adams et al., 2020). This presence of RBBP4 in these SIN3A/B complexes is not surprising since RBBP4 is a known component of the canonical SIN3-HDAC complex (Hassig et al., 1997). Interestingly, when Adams and colleagues did the reverse experiment and immunoprecipitated RBBP4, they readily detected SIN3A/B, but also a strong enrichment of MuvB proteins (Adams et al., 2020), which suggests that at least in this experimental setup, RBBP4 exclusively binds to either MuvB or SIN3. The conflicting results gained from IP/mass spec experiments are reflected by several studies that analyzed interactions of SIN3 and MuvB proteins by IP/Western. While we and others (Adams et al., 2020; Litovchick et al., 2007; Silverstein and Ekwall, 2005; van Oevelen et al., 2008) have been unable to coprecipitate endogenous SIN3B/HDAC with DREAM complex components, two publications showed binding of SIN3B and HDAC1 to DREAM (Bainor et al., 2018; Pilkinton et al., 2007).

Here we aimed to identify more conclusively the role of SIN3 and HDACs in DREAM function and more broadly in repression of cell-cycle genes. Previous genetic knockout of SIN3B increased the expression of about 100 DREAM target genes in serum-starved T98G cells, and based on the observation that H3K9 acetylation was increased at the promoters of *CCNA2* and *INCENP*, it was suggested that SIN3B mediates the repressive function of DREAM through its ability to tether histone modifiers to chromatin (Bainor et al., 2018). In contrast, we found that loss of SIN3B does not derepress cell-cycle genes in HCT116 and C2C12 cells arrested by activation of the p53 pathway or serum starvation (Figs. 2, 4, 5). We reproduced observations of a defect in cell-cycle gene repression in SIN3B^-/-^ T98G cells upon serum starvation, however CDK4/6 inhibition with Palbociclib rescued the knockout phenotype (Fig. 3), which suggests that SIN3B functions upstream of DREAM in this context. In contrast, the loss of DREAM repressor function we found previously in arrested LIN37^-/-^ NIH3T3, HCT116, and C2C12 cells (Mages et al., 2017; Uxa et al., 2019) persisted in serum-starved and Palbociclib-treated LIN37^-/-^ T98G cells (Fig. 3). Despite these diverging results regarding functional interactions between DREAM and SIN3:HDAC complexes, consistent data have been published showing that SIN3A, SIN3B, and HDAC1/2 bind to G1/S and G2/M cell-cycle gene promoters (Bainor et al., 2018; Rayman et al., 2002; van Oevelen et al., 2010; van Oevelen et al., 2008). These data are in line with our *in silico* analysis (Fig. 1) and ChIP-qPCR results from arrested HCT116 cells (Fig. 2G). In differentiated C2C12 cells, tiling array data showed that E2F4 binding to promoters is centered around the TSS, while peaks of SIN3A and SIN3B are shifted about 200bp downstream of the TSS (van Oevelen et al., 2008). Even though the authors suggested that E2F4 and SIN3A/B were recruited as a complex, the mechanistic details of a potential E2F-dependent recruitment of SIN3 followed by spreading to downstream regions remained unclear. While the authors did not analyze the effect of single SIN3A or SIN3B knockdown in this cellular system, combined knockdown of SIN3A and SIN3B resulted in an about 3-fold upregulation of several cell-cycle genes (*Brca1, Top2a, Mcm5, Kif20a, Aurkb*), which reflects the level of derepression we find in SIN3A/B-depleted arrest HCT116 cells (Fig. 5). Based on these and our results, it is likely that both SIN3A and SIN3B bind to a broad set of cell-cycle genes, can substitute for each other, and generally contribute to the repression of cell-cycle repression.

It remains to be elucidated how SIN3 proteins get recruited to cell-cycle gene promoters and how they contribute to repression, given that they neither contain a DNA-binding domain nor enzymatic activity (Chaubal and Pile, 2018). Our results suggest that SIN3 proteins function at cell-cycle gene promoters independent of DREAM and RB, since we find a general increase of derepression in arrested cells depleted of LIN37/SIN3A/SIN3B or RB/SIN3A/SIN3B (Fig. 5E, Suppl. Fig. 1E). Since DREAM still forms and binds to target genes in LIN37-negative cells (Mages et al., 2017; Uxa et al., 2019), the complex could potentially still be able to recruit SIN3 proteins, which could contribute to target gene repression. However, we have shown that loss of LIN37 derepresses cell-cycle gene promoters equally as strongly as loss of DREAM-binding (Mages et al., 2017), and another study showed that binding of SIN3B to several cell-cycle gene promoters persisted in p130^-/-^;p107^-/-^ and Rb^-/-^ serum-starved MEFs (Rayman et al., 2002). Based on these results, it appears unlikely that SIN3 proteins get recruited to cell-cycle genes via DREAM or RB. Since HDACi did not phenocopy loss of SIN3A/B (Fig. 5, 6, Suppl. Fig. 1, 2), it is also likely that the observed SIN3-dependent effects do not rely on recruitment of HDACs, but on one of the many chromatin-modifying enzymes or transcription factors that have been described as interactors of SIN3 proteins (Chaubal and Pile, 2018).

Many studies have analyzed the effects of HDACi on transformed and non-transformed proliferating cells, and generally, treatment with HDAC inhibitors increases the expression of anti-proliferative and pro-apoptotic genes, represses the expression of pro-proliferative genes, and results in cell-cycle arrest and apoptosis (Bolden et al., 2013; Chan et al., 2013; Han et al., 2000; Laporte et al., 2017; Mazzio and Soliman, 2018; Moreira et al., 2003; Peart et al., 2005; Roy et al., 2008; Vinodhkumar et al., 2008; Wang et al., 2015). These results can be recapitulated by combined knockout of HDAC1 and HDAC2 (Yamaguchi et al., 2010). Since cancer cells are generally more sensitive to HDACi than non-transformed cells, HDAC inhibitors are promising drugs in cancer therapy with four molecules approved by the FDA, and a multitude of ongoing clinical trials (Ramaiah et al., 2021). A central mechanism that initiates cell-cycle arrest and apoptosis following the loss of HDAC1/2 activity is an increased expression of the CDK inhibitors p21, p27, and p57 (Richon et al., 2000; Yamaguchi et al., 2010). p21 expression is further stimulated through acetylation and stabilization of p53 (Ramaiah et al., 2021). Reduced CDK activity from high inhibitor levels results in the accumulation of unphosphorylated pocket proteins followed by formation of E2F:RB and DREAM complexes. We and others have shown that these complexes are essential for inducing G0/G1 arrest (Dannenberg et al., 2000; Mages et al., 2017; Sage et al., 2000; Schade et al., 2019b; Uxa et al., 2019). Therefore, if HDAC activity is necessary for the repression of cell-cycle genes by RB and DREAM, as several studies have shown (Bainor et al., 2018; Brehm et al., 1998; Ferreira et al., 1998; Ferreira et al., 2001; Lai et al., 1999; Luo et al., 1998; Magnaghi-Jaulin et al., 1998; Siddiqui et al., 2003; Zhang et al., 2000), how could it be that HDACi results in arrest of cancer cells? A possible explanation of this paradox could be that E2F:RB and DREAM can partially repress their target genes without recruiting HDAC activity, but HDACs are essential to completely shut down cell-cycle gene transcription. In this context, gene expression data originating from proliferating cells treated with HDAC inhibitors are not particularly helpful since they do not provide information on whether genes are completely repressed upon treatment. Thus, we addressed this question by inducing strong repression of cell-cycle genes through activation of the p53-p21 pathway, and then added HDAC inhibitors to analyze if repression is relieved. Even though we found that HDACi increases histone tail acetylation at cell-cycle gene promoters (Fig. 6A), we did not detect a global derepression in sets of representative G1/S and G2/M genes (Fig. 6, Suppl. Fig. 2).

Our results are another example that suggests that histone marks previously associated with actively expressed genes do not directly cause or even correlate with transcriptional activation (Morgan and Shilatifard, 2020). Many of the original data connecting pocket-protein-dependent repression to HDAC activity were generated with *in vitro* approaches, reporter assays, and over-expressed proteins (Bainor et al., 2018; Brehm et al., 1998; Ferreira et al., 1998; Ferreira et al., 2001; Lai et al., 1999; Luo et al., 1998; Magnaghi-Jaulin et al., 1998; Siddiqui et al., 2003; Zhang et al., 2000). Using such artificial methods may have led to an overinterpretation of the results, and slight changes found in the mRNA expression of a few tested genes were later generalized to the regulation of E2F-dependent genes. In contrast, our data suggest that HDAC activity is not generally required for cell-cycle gene repression during reversible cell-cycle arrest and that alternative mechanisms like RB-dependent inhibition of activator E2Fs (Helin et al., 1993; Hiebert et al., 1992), stabilizing nucleosomes by DREAM (Asthana et al., 2022), and potentially recruitment of other chromatin-modifying or nucleosome-remodeling proteins are sufficient to induce robust repression. However, HDAC activity may be involved in fine-tuning the activity of subsets of cell-cycle genes under specific physiological conditions (Sanidas et al., 2019). Furthermore, in contrast to reversible cell-cycle arrest, HDAC activity may be required for the permanent silencing of cell-cycle genes during terminal cell-cycle exit. This topic has yet to be well-studied, but several transcriptome data sets derived from terminally differentiated cells negative for HDAC1/2, or treated with HDAC inhibitors, are available and suggest that HDAC activity is also not generally required in this context (Montgomery et al., 2007; Rai et al., 2008). Taken together, it remains to be elucidated in which biological contexts SIN3 proteins and HDAC activity substantially contribute to cell-cycle gene regulation.

## Methods

### Cell culture and drug treatment

HCT116, T98G, C2C12, A549, and BJ-hTert wild-type and knockout cells were grown in Dulbecco’s modified Eagle’s medium (Gibco, #10569044) supplemented with 10 % fetal calf serum (Corning, #MT35010CV) and penicillin/streptomycin (Gibco, #15140122). Cells were maintained at 37°C and 10% CO_2_ and were tested negative for mycoplasma contamination by PCR with a mixture of primers that have been described previously (Uphoff and Drexler, 2002). For induction of p53, cells were treated with Idasanutlin (5 μM; R&D Systems, #12-35-22-07) or Doxorubicin (0.5 µM; Selleckchem, #E2516). T98G cells were starved in DMEM containing 0% FBS, and C2C12 cells were starved with DMEM containing 0.1% FBS. CDK4/6 inhibition was performed with Palbociclib (10 µM, Selleckchem, #S4482). Histone deacetylases were inhibited with Romidepsin (4 nM; Active Motif, # 14083) or Panobinostat (20 nM; Selleckchem, # S1030).

### Generation of knockout cell lines by CRISPR/Cas9 nickase

*SIN3B^-/-^, SIN3B^-/-^;LIN37^-/-^*, *SIN3B^-/-^;RB^-/-^* HCT116 cells, *SIN3B^-/-^*, *LIN37^-/-^*, *SIN3B^-/-^;LIN37^-/-^* T98G cells, and *SIN3B^-/-^*C2C12 cells were created by CRISPR/Cas9 nickase, applying the pX335-U6-Chimeric_BB-CBh-hSpCas9n(D10A) vector (Ran et al., 2013a; Ran et al., 2013b) as described earlier (Mages et al., 2017; Uxa et al., 2019). Mutations were introduced into exons 3 or 4 of the human SIN3B gene, exon 5 of the mouse Sin3b gene, and exon 6 of the LIN37 gene. Sequences of the oligonucleotides are provided in Supplementary Table S1. Knockout was confirmed by SDS-PAGE and Western blot.

### RNA interference

HCT116 cells were cultivated in 6-well plates and transfected with 20 nM SIN3A siRNAs (Horizon Discovery, #MQ-012990-00-0002) or a non-targeting control siRNA (siGENOME Non-Targeting siRNA 5, Horizon Discovery, # D-001210-05-05) and 5 µl Lipofectamine RNAiMAX (Invitrogen, #13778075) in a total volume of 2 ml in antibiotics free 10% DMEM. 24 h after transfection, cells were treated with either DMSO or 5 µM Idasanutlin (R&D Systems, #12-35-22-07) for an additional 24 h before cells were harvested for protein and RNA extraction.

### RNA extraction, reverse transcription, and semi-quantitative real-time PCR

Total RNA was isolated with the Direct-zol RNA MiniPrep Kit (Zymo Research, #R2053). One-step RT-qPCR was performed using the GoTaq 1-Step RT-qPCR System (Promega, #A6020) and MicroAmp Fast Optical 96 Well Reaction Plates (Applied Biosystems, #4346907) on a Quantstudio 3 Real-Time PCR cycler (ThermoFisher Scientific). See Supplementary Table S1 for primer sequences.

### SDS-PAGE and Western blot

Whole-cell extracts prepared with RIPA buffer (10 mM Tris-HCl pH 8.0, 150 mM NaCl, 1 mM EDTA, 0.1% SDS, 0.1% Sodium Deoxycholate, 0.1%TritonX) or immunoprecipitated samples were analyzed by SDS-PAGE and Western blot following standard protocols as described earlier (Kirschner et al., 2008). See Supplementary Table S2 for the list of antibodies used for Western blotting.

### Chromatin immunoprecipitation (ChIP)

Cells were harvested, cross-linked with PBS (Gibco, # 14190250) supplemented with 1% paraformaldehyde (Electron Microscopy Sciences, #50-980-487), and quenched with 125 mM Glycine (Fisher Scientific, # BP3815). Nuclei were isolated using Buffer A (Cell Signaling, #7006S) and Buffer B (Cell Signaling. #7007S). MNase enzyme was prepared in-house using Addgene plasmid # 136291 (Yan et al., 2019). Nuclei were MNase-treated on ice for 30 minutes followed by 15 minutes of incubation at 37°C and 5x direct sonication for 1s to create ∼300bp chromatin fragments. Protein-DNA complexes were immunoprecipitated with the indicated antibodies overnight at 4°C and bound to Pierce protein A/G magnetic beads (ThermoScientific, #PI88803). Beads were subsequently washed with the following buffer types in order: 6x RIPA (10 mM Tris-HCl pH 8.0, 1 mM EDTA, 0.1% SDS, 0.1% Sodium Deoxycholate, 0.1%TritonX) supplemented with 140 mM NaCl, 3x RIPA supplemented with 500 mM NaCl, 3x LiCl buffer (10 mM Tris-HCl pH 8.0, 250 mM LiCl, 1 mM EDTA, 0.5% Sodium Deoxycholate, 0.5% Triton X), and 3x 10 mM Tris-HCl 8.0 (salt-free). Precipitants were eluted twice with 150 µl elution buffer (10 mM Tris-HCl 8.0, 5 mM EDTA, 300 mM NaCl, 0.6% SDS) with 30s vortexing and 15 min incubation at 37°C. Eluants were treated with RNaseA (Thermo Scientific, #FEREN0531) for 30 minutes at 37°C, then treated with Proteinase K (Thermo Scientific, #EO0491) for 1 hour at 55°C, and reverse cross-linked at 65°C overnight. DNA was purified using Zymo DNA Clean & Concentrator-5 kits (Zymo Research, #77001-152). qPCR was performed with the GoTaq qPCR Master Mix (Promega, #A6001) on a Quantstudio 3 Real-Time PCR System (ThermoFisher Scientific). See Supplementary Table S1 for primer sequences and Supplementary Table S2 for antibodies.

### EdU assay

EdU assays were performed with the Click-iT^®^ EdU Flow Cytometry Assay Kit (ThermoFisher Scientific, #C10420) according to the manufacturer’s information. 10,000 cells of each sample were analyzed by flow cytometry (LSR II, BC). Data were analyzed with FlowJo (BD).

### HDAC activity assay

Cells were lysed in lysis buffer (50 mM Tris pH 8.0, 10 mM MgCl2, 0.2% Triton X, 300 mM NaCl) by 5x direct sonication for 1s. Lysates were clarified by centrifugation (13,000 rpm, 10 min, 4°C), and NaCl concentration was adjusted to 150 mM. 2-3 µg of antibodies were incubated with Pierce protein A/G magnetic beads (ThermoScientific, #PI88803) for 1 h on a rotator at 4°C. Beads were washed 3x with the buffer previously described with 150 mM NaCl. 5mg protein extracts were incubated with the antibody-bound beads for 3 h on a rotator at 4°C. Beads were washed 5x with lysis buffer containing 150 mM NaCl and resuspended in 500 µl HDAC-Glo buffer (Promega, #G648A). Beads were serially diluted with HDAC-Glo buffer, and 20µl of each dilution was transferred into a 384-well, white, flat-bottom plate. HDAC activity of three technical replicates was measured using the HDAC-Glo I/II Assay (Promega, #G6420) following the manufacturer’s protocol on-bead with an Envision plate reader (Perkin Elmer). The remaining beads were boiled in Laemmli buffer (Laemmli, 1970) and subjected to SDS-PAGE and Western blot analysis.

### Next generation sequencing and transcriptome analysis

Total RNA was isolated from HCT116 cells with the Direct-zol RNA MiniPrep Kit (Zymo Research, #R2053). Library preparation, rRNA depletion, and Illumina sequencing were performed at Genewiz/Azenta Life Sciences. Reads were trimmed with trimgalore (version 0.6.10, https://www.bioinformatics.babraham.ac.uk/projects/trim_galore/) using cutadapt version 4.2 (Martin, 2011) and fastqc v0.12.1 (http://www.bioinformatics.babraham.ac.uk/projects/fastqc) for quality control. GNU parallel was used for parallelization (Tange, 2022). After trimming, 0.5 % to 0.7% of reads were too short to be considered mappable (<20 nt). Trimmed reads were mapped to the hg38 genome using segemehl (version 0.3.4) (Hoffmann et al., 2014) using standard parameters and the -S option to be able to map spliced reads. 42 to 53 million reads were mapped per sample (93 % to 96 % of all reads). Between 86 % and 91 % of all reads were mapped uniquely. The mapped reads were annotated using featureCounts (Liao et al., 2014) version 2.0.3 against the gencode v.27 annotation, using the following parameters: -p -t exon -g gene_id. The resulting reads per gene counts were normalized and analyzed using DESeq2 (Love et al., 2014) to find differentially expressed genes. Comparisons of conditions and normalizations were done pairwise. All reported and plotted p-values are multiple hypotheses adjusted by DESeq2 with the Benjamini and Hochberg method. To calculate enrichment of LIN37 targets among up-regulated genes (Fig. 5a) we used the hypergeometric test as implemented in the phyper function in the stats package in R (R_Core_Team, 2021). GO term analyzes were performed with ShinyGo (Ge et al., 2020), and Venn diagrams were built with Venny (https://bioinfogp.cnb.csic.es/tools/venny/index.html).

## Availability

RNA-Seq data generated for this publication was deposited at GEO GSE240734.

## Acknowledgement

Technical support was provided by Beverley Rabbitts, UCSC Chemical Screening Center, RRID SCR_021114.

## Author contributions

G.A.M. and S.M.R. conceived and supervised the study. A.R. evaluated the RNA-seq data. A.B., M.R.S, G.A.M., T.U.W, and K.M.R. acquired all other data. G.A.M., S.M.R., A.B., and A.R. wrote the original draft of the manuscript. M.R.S., T.U.W, and K.M.R. reviewed and edited the manuscript.

## Supplementary Data

**Suppl. Fig. 1**

**Suppl. Fig. 2**

**Suppl. Table S1:** Sequences of oligonucleotides used in this study.

**Suppl. Table S2:** Antibodies used in this study.

**Suppl. Table S3:** G1/S, G2/M, and non-DREAM/non-cell cycle regulated control genes selected to generate Fig. 1.

**Suppl. Table S4:** Differentially expressed genes in SIN3B^-/-^, SIN3A knockdown, and SIN3B^-/-^;SIN3A knockdown HCT116 cells.

