## Supplemental Figures for "HDAC activity is dispensable for repression of cell-cycle genes by DREAM and E2F:RB complexes"

### Supplementary Figures

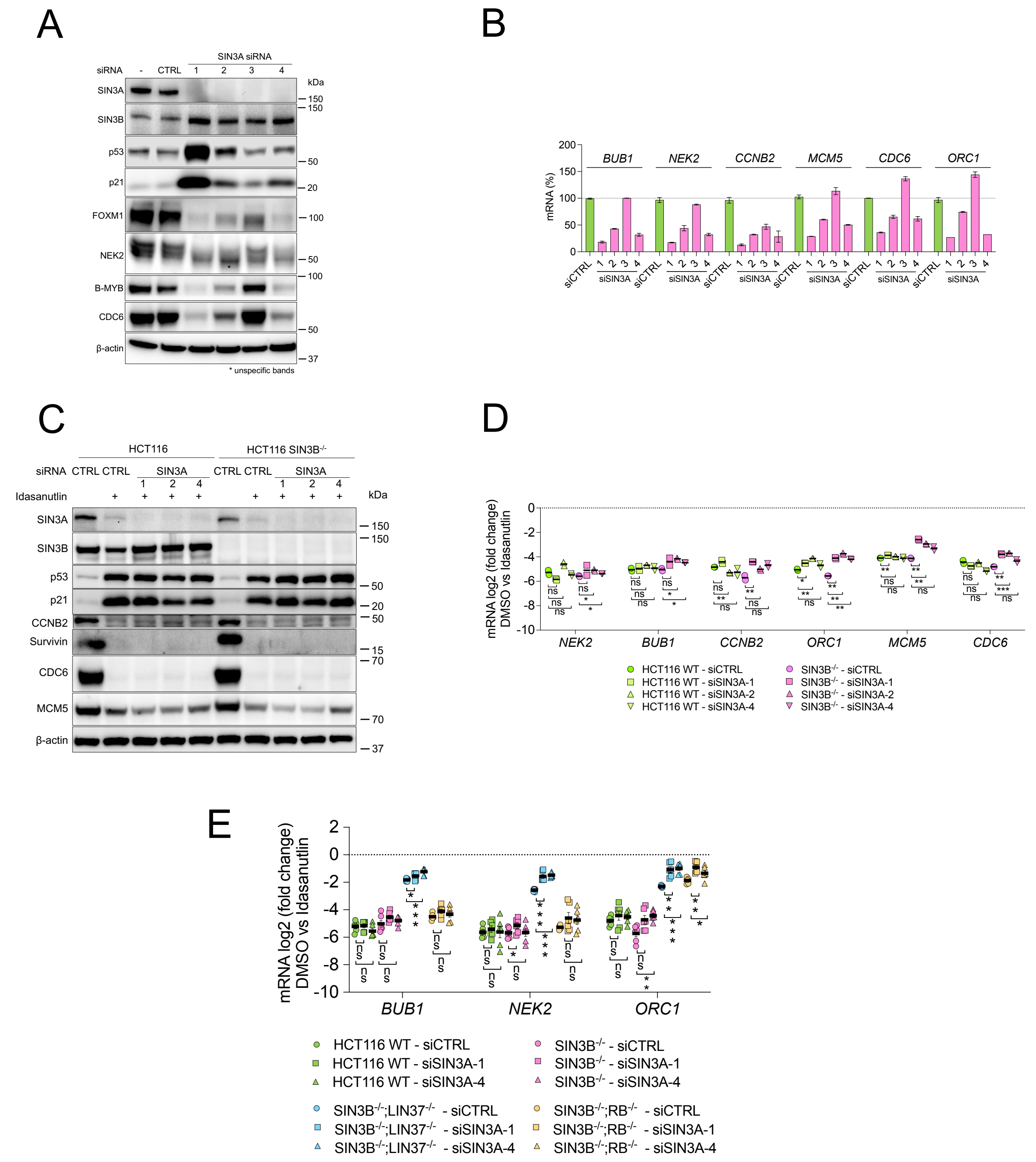


**Suppl. Fig. 1: Combined loss of SIN3A and SIN3B increases cell-cycle gene mRNA expression in arrested HCT116 cells, while loss of SIN3A alone has only minor effects.** (A) HCT116 cells were transfected with four SIN3A siRNAs or a non-targeting control siRNA (CTRL), and protein expression was analyzed after 48 h by Western blotting. Mean values of a representative experiment (two experiments total) with two technical replicates (± SD) are shown. (B) mRNA expression of representative cell-cycle genes in cells treated as described in (A) was analyzed by RT-qPCR (one representative experiment, two experiments total, mean values ± SD). (C) HCT116 wild-type and SIN3B$^{-/-}$ cells were transfected for 48 h with three SIN3A-targeting siRNAs or a non-targeting control siRNA, and 5 µM Idasanutlin was applied for the final 24 h. Protein levels were analyzed by Western blotting, and (D) mRNA expression of cell-cycle genes was measured by RT-qPCR. One representative experiment out of two replicates with two technical replicates is shown. (E) mRNA expression of cell-cycle genes that were not identified as significantly upregulated by RNA-seq in arrested HCT116 cells depleted of SIN3A and SIN3B was analyzed by RT-qPCR. Averages of three biological and two technical replicates are given (mean values ± SEM). Significances were calculated with the two-tailed Student's T-Test (ns – not significant, * p ≤.05, ** p ≤ .01, *** p ≤ .001).

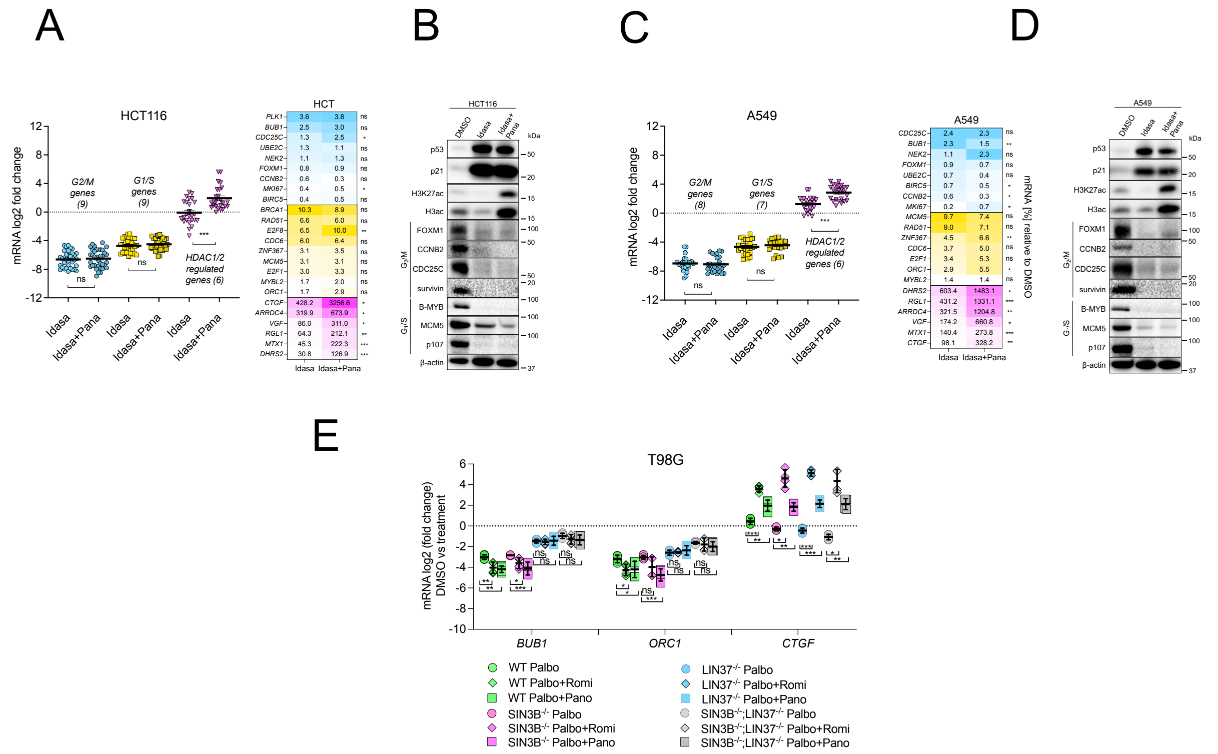


**Suppl. Fig. 2: HDACi by the pan-HDAC inhibitor Panobinostat does not lead to a general upregulation of DREAM and E2F:RB target genes in arrested cells.** (A, B) HCT116 and (C, D) A549 were treated with 5 µM Idasanutlin for 48 h and 20 nM Panobinostat for the final 24 h. (A, C) mRNA expression was evaluated by RT-qPCR and compared to DMSO-treated cells for each respective line. Expression data are shown as gene-set clusters (left) and individual genes (right). The datasets contain two biological replicates with two technical each presented as mean values ± SEM. (B, D) Protein expression and histone acetylation were evaluated via Western blotting with the indicated antibodies. A biological replicate of each Western blot experiment produced similar results. (E) The indicated T98G wild-type and knockout lines (two clones each) were treated with 10 µM Palbociclib for 48 h and with 4 nM Romidepsin or 20 nM Panobinostat for the final 24 h when indicated. mRNA levels were evaluated by RT-qPCR (two technical replicates) and compared to DMSO-treated cells. Mean values ± SD are shown. Significances in (A), (C), and (E) were calculated with the two-tailed Students T-Test (ns – not significant, * p ≤.05, ** p ≤ .01, *** p ≤ .001).
